# A zebrafish model of COVID-19-associated cytokine storm syndrome reveals that the Spike protein signals via TLR2

**DOI:** 10.1101/2022.07.14.500031

**Authors:** Sylwia D. Tyrkalska, Alicia Martínez-López, Annamaria Pedoto, Sergio Candel, María L. Cayuela, Victoriano Mulero

## Abstract

Understanding the mechanism of virulence of SARS-CoV-2 and host innate immune responses are essential to develop novel therapies. One of the most studied defense mechanisms against invading pathogens, including viruses, are Toll-like receptors (TLRs). Among them, TLR3, TLR7, TLR8 and TLR9 detect different forms of viral nucleic acids in endosomal compartments, whereas TLR2 and TLR4 recognize viral structural and nonstructural proteins outside the cell. Although many different TLRs have been shown to be involved in SARS-CoV-2 infection and detection of different structural proteins, most studies have been performed *in vitro* and the results obtained are rather contradictory. In this study, we report using the unique advantages of the zebrafish model for *in vivo* imaging and gene editing that the S1 domain of the Spike protein from the Wuhan strain (S1WT) induced hyperinflammation in zebrafish larvae via a Tlr2/Myd88 signaling pathway and independently of interleukin-1β production. In addition, S1WT also triggered emergency myelopoiesis, but in this case through a Tlr2/Myd88-independent signaling pathway. These results shed light on the mechanisms involved in the COVID-19-associated cytokine storm syndrome.

## INTRODUCTION

Since the onset of the COVID-19 pandemic that began in late 2019, research on severe acute respiratory syndrome coronavirus 2 (SARS-CoV-2) and its disease has been conducted extensively in many different directions. Viral infections are very complex processes and require research at different interdisciplinary levels to obtain information on basic pathways leading to detailed explanations of the pathogenicity of the virus, host immune response and treatment development. Toll-like receptors (TLRs) are evolutionarily conserved pattern recognition receptors (PRRs) that discriminate between self and non-self by detecting pathogen-associated molecular patterns (PAMPs) and initiate the immune response activating signaling cascades that lead to the production of antimicrobial and proinflammatory molecules [1,2]. All TLRs belong to type I transmembrane proteins, composed of an amino-terminal leucine-rich repeat-containing ectodomain (responsible for PAMP recognition), a transmembrane domain and cytoplasmic carboxy-terminal Toll-interleukin-1 receptor (IL-1R) homology (TIR) domain (responsible for activation of downstream signal transduction) [3], which is also present in the interleukin-1 receptor (IL-1R). TLRs signals via the myeloid differentiation primary response 88 (MYD88) or TIR-domain containing adaptor inducing interferon-β (TRIF) [4,5]. Importantly, MYD88-dependent pathway can be activated by all TLRs except TLR3, which only signals through TRIF, while TLR4 activates both pathways [6,7].

To date there are many well characterized TLRs that have been linked to antiviral immunity. Among them, TLR3, TLR7, TLR8 and TLR9 detect different forms of viral nucleic acids in endosomal compartments, while TLR2 and TLR4 are able to recognize viral structural and nonstructural proteins outside the cell [8,9]. TLR2 is located on the plasma membranes of immune, endothelial, and epithelial cells [10] to recognize mainly components of microbial cell walls and membranes, such as lipoproteins and peptidoglycans. As a heterodimeric receptor, it is paired with TLR1 or TLR6 to recognize different bacterial products such as triacylated lipopeptides (TLR1/TLR2) and diacylated lipopeptides (TLR1/TLR6). In case of viruses, it is assumed that TLR2 is able to recognize enveloped viral particles [11].

It has been shown that many different TLRs may be involved in the SARS-CoV-2 infection. Both extracellular and intracellular TLR family receptors have been shown to play a role in SARS-CoV-2 viral detection. TLR2 recognizes the SARS-CoV-2 envelope protein, resulting in MYD88-dependent inflammation [12]. TLR3 is presumed to be critical in the recognition of double stranded RNA (dsRNA) from SARS-CoV-2, which is generated during viral replication, and stimulates endosomal TLR3 in addition to other intracellular receptors [13]. TLR4 signaling by MYD88 and TRIF dependent pathways is proposed to detect viral structural proteins and glycolipids [14,15]. Finally, TLR7 has been linked to COVID-19 severity in multiple studies, strongly suggesting a key role for TLR7 in COVID-19 pathogenesis [13].

Although TLRs are highly conserved during evolution, in fish they show different characteristics than those present in mammals [16]. In zebrafish, high expression of TLRs was detected in the skin, which may suggest their important role in the defense against pathogens. It is also worth mentioning that zebrafish has an almost complete set of 20 putative TLR variants, of which 10 have direct human orthologs, including TLR2 [17,18]. TLR22 belongs to a fish-specific subfamily and recognizes double stranded RNA [19], whereas TLR21 is present in birds, amphibians and fish with similar expression profiles and activity to TLR9 [20]. Moreover, zebrafish show duplication of some mammalian TLRs including Tlr4ba/Tlr4bb for TLR4 and Tlr5a/Tlr5b for TLR5, and Tlr8a/Tlr8b for TLR8 [21]. Interestingly, homologs of mammalian TLR6 and TLR10 are absent in fish, while other TLRs, such as Tlr14 and Tlr18, have also been identified. In addition, ligands of some zebrafish Tlr receptors have already been identified, such as lipoproteins, lipopeptides or Pam3CSK4 that can be recognized by the heterodimers of Tlr2 and flagellin by Tlr5 [22]. Furthermore, the TLR/IL-1R downstream signaling pathway is also highly conserved in zebrafish including the ortholog of Myd88 among others [23].

In this study, we show using the zebrafish model that the hyperinflammation induced by the Spike protein of SARS-CoV-2 is mediated via a Tlr2/Myd88 signaling pathway and independently of Il1b production. In addition, Spike protein-induced emergency myelopoiesis is independent of Tlr2/Myd88 in this model.

## RESULTS

### Myd88 is required for hyperinflammation but dispensable for emergency myelopoiesis induced by S1WT

As the specific TLRs activated by the spike protein of SARS-CoV-2 are controversial, either recombinant S1 from the Wuhan strain (S1WT) or flagellin (positive control that activates Tlr5) were injected into the hindbrain of Myd88-deficient zebrafish and the recruitment to the injection site and the total number of neutrophils and macrophages in the head and the whole body were analyzed at 6, 12 and 24 hours post-injection (hpi). Although robust recruitment of both neutrophils and macrophages was observed in wild type larvae, a significantly lower recruitment of both immune cells was observed in Myd88-deficient larvae at 6 and 12 hpi (Figures 1A and 1B). In contrast, no differences in total number of neutrophils and macrophages in the head or whole body were observed between mutant and wild type larvae at any timepoint (Figure 1A and 1B). These results suggest that S1WT-induced emergency myelopoiesis is Myd88-independent and, therefore, depends exclusively on the inflammasome [24].

**Figure 1:**
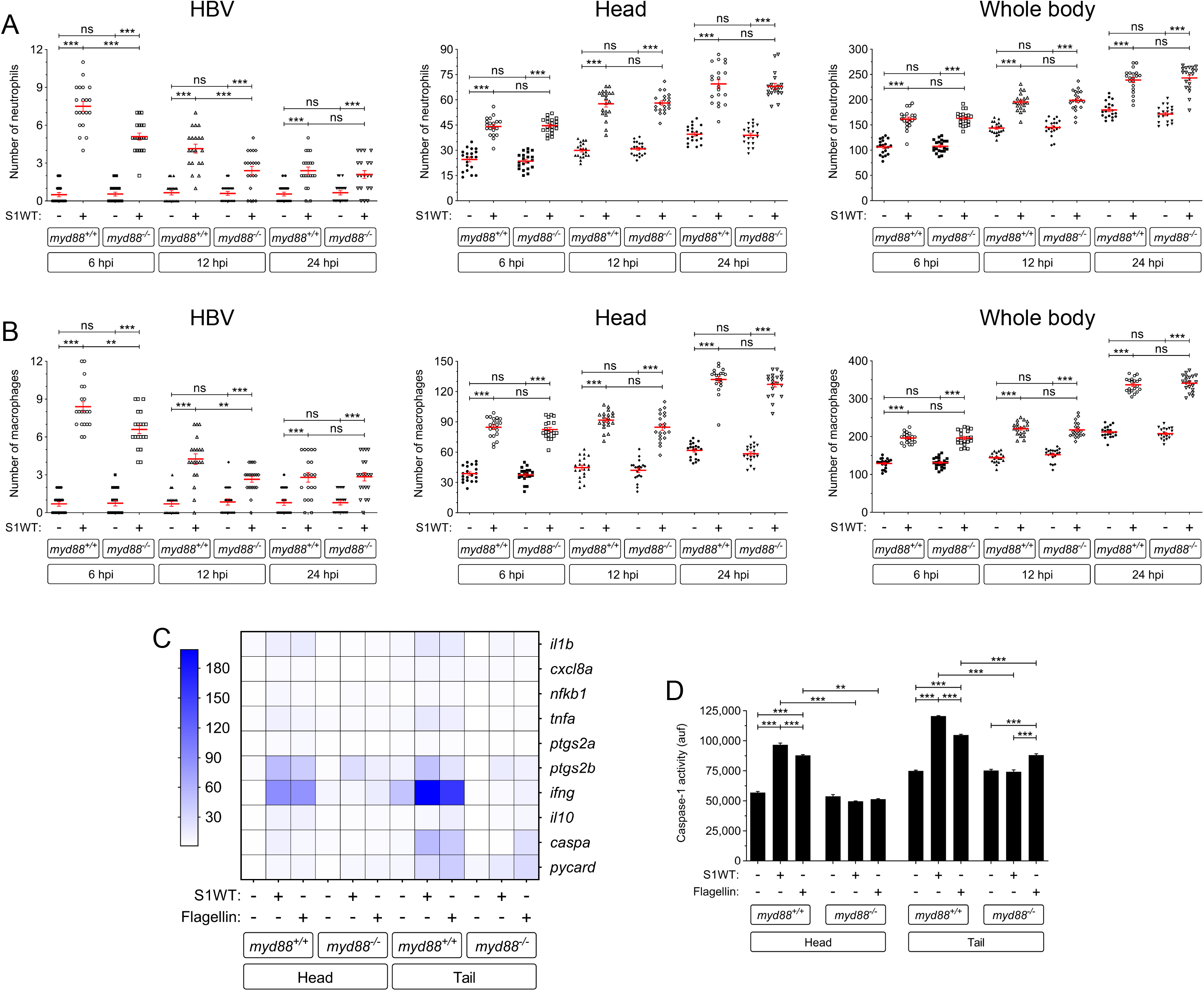
Myd88 is required for hyperinflammation but dispensable for emergency myelopoiesis induced by S1WT. Recombinant S1WT (+) or vehicle (-) were injected in the hindbrain ventricle (HBV) of 2 dpf wild type and Myd88-deficient *Tg(mpx:eGFP)* (A) or wild type (B-D) larvae. Neutrophil (A) and macrophage (neutral red positive cells) (B) recruitment and number were analyzed at 6, 12 and 24 hpi by fluorescence (A) or brightfield (B) microscopy, the transcript levels of the indicated genes (D) were analyzed at 12 hpi by RT-qPCR in larval head and tail (E), and caspase-1 activity was determined at 24 hpi using a fluorogenic substrate (F). Each dot represents one individual and the mean ± S.E.M. for each group is also shown. P values were calculated using one-way ANOVA and Tukey multiple range test. RT-qPCR data are depicted as a heat map in D with higher expression shown in darker color. ns, not significant, **p≤0.01, ***p≤0.001. auf, arbitrary units of fluorescence.

To further analyze the impact of Myd88 in the local and systemic inflammation induced by S1WT, samples were collected at 12 hpi from heads and the rest of the body for RT-qPCR analysis. Although flagellin and S1WT induced similar gene expression patterns, Myd88 deficiency impaired the induction of transcript levels of genes encoding inflammatory mediators Il1b, Cxcl8a, Nfkb1, Tnfa, Ptgs2a, Ptgs2b, Infg and Il10 by both S1WT and flagellin (Figures 1C and S1A–S1H). Curiously, mRNA levels of genes encoding the canonical inflammasome effector Caspa also appear to be Myd88 dependent, as S1WT failed to induce their expression in Myd88-deficient larvae, while transcript levels of the gene encoding the inflammasome adaptor Asc (*pycard* gene) were largely unaffected (Figures 1C, S1I and S1J). Similarly, Myd88 deficiency impaired the induction of caspase-1 activity by S1WT and flagellin locally and systemically. These results suggest that Myd88 plays a key role in the hyperinflammation induced by S1WT in zebrafish.

### Il1b signaling is not involved in S1WT-induced hyperinflammation in zebrafish

To learn if the hyperinflammation induced by S1WT was dependent of Tlr signaling, we knocked down Il1b, as its receptor also signals through Myd88. For this purpose, we used a specific guide RNA that provided 70% of efficiency and showed neither toxicity nor malformations in embryos (Figures S2A and S3A–S3C). The results showed that Il1b-deficiency failed to affect S1WT-induced neutrophil and macrophage recruitment, neutrophilia and monocytosis (Figures 2A and 2B). In addition, transcript levels of genes encoding major inflammatory molecules were similarly induced by S1WT in wild type and Il1b-deficient larvae, apart from those of *il1b* itself which were drastically induced and those of *ptgs2a* and *ptgs2b* which were weakly decreased (Figures 2C and S4A-S4F). Curiously, *i110* and *infg* mRNA levels were systemically higher in Il1b-deficient larvae than in their wild type siblings (Figures 2C, S4G and S4H), while those of genes encoding the inflammasome components Caspa and Pycard, and caspase-1 activity were rather similar in Il1b-deficient and wild type larvae (Figures 2C, 2E, S4I and S4J). These results taken together confirmed the impact of the genetic edition of *il1b* gene and that S1WT-induced hyperinflammation is largely Il1b-independent.

**Figure 2:**
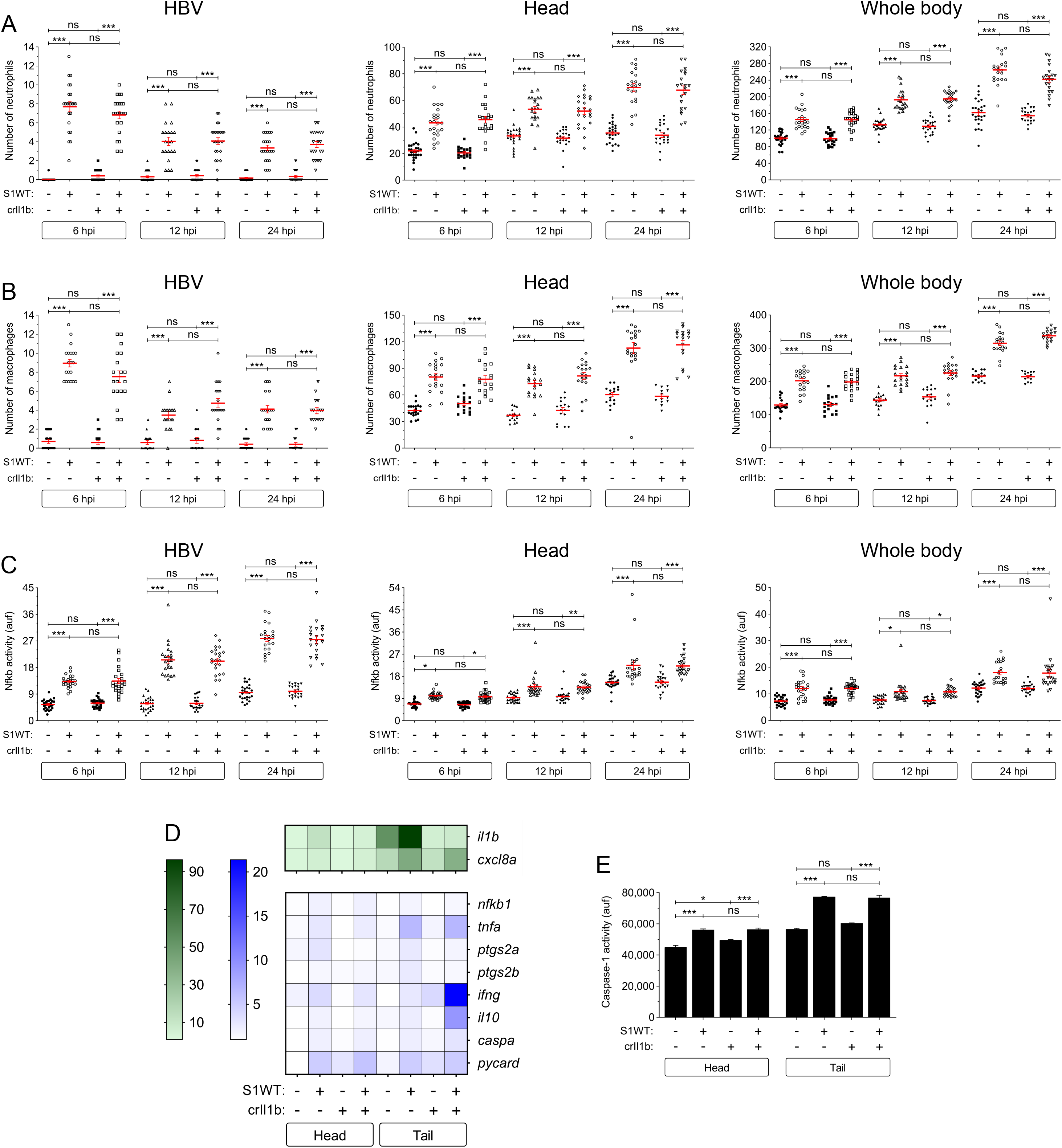
Il1b signaling is not involved in S1WT-induced hyperinflammation in zebrafish. One-cell stage zebrafish eggs of *Tg(lyz:dsRED2)* (A), *Tg(mfap4:mCherry)* (B), *Tg(NFkB-RE:eGFP)* (C) and wild type (D, E) were microinjected with control or *il1b* crRNA/Cas9 complexes. At 2 dpf, recombinant S1WT (+) or vehicle (-) were injected in the hindbrain ventricle (HBV) of control and Il1b-deficient larvae. Neutrophil (A) and macrophage (B) recruitment and number, and Nfkb activation (C) were analyzed at 6, 12 and 24 hpi by fluorescence microscopy, the transcript levels of the indicated genes were analyzed at 12 hpi by RT-qPCR (D), and caspase-1 activity was determined at 24 hpi using a fluorogenic substrate (E). Each dot represents one individual and the mean ± S.E.M. for each group is also shown. RT-qPCR data are depicted as a heat map in D with higher expression shown in darker color. P values were calculated using one-way ANOVA and Tukey multiple range test. ns, not significant, *≤p0.05, **p≤0.01, ***p≤0.001. auf, arbitrary units of fluorescence.

### Tlr2 mediates the S1WT-induced hyperinflammation in zebrafish

Since Myd88 acts downstream of almost all TLRs apart from TLR3 [6,7], we decided to check the expression levels of the orthologs of the TLRs shown to be involved in the responses to SARS-CoV-2, namely *tlr2, tlr4ba, tlr4bb* and *tlr7*, and *tlr3* as a negative control, upon S1WT hindbrain injection. Notably, only the transcript levels of *tlr2* increased after S1WT injection (Figures S5A-E), suggesting S1WT signals via Tlr2 in zebrafish. We then knocked down Tlr2 using a specific guide RNA that resulted in approximately 70% efficiency (Figure S2B) and showed no detrimental effect in larval development (Figures S3D–S3F). We found that Tlr2 deficiency did not affect S1WT-induced neutrophilia and monocytosis at any of the times tested (Figures S3A and 3B), further confirming that S1WT-induced emergency myelopoiesis is Tlr2/Myd88 independent. However, neutrophil and macrophage recruitment at the S1WT injection site was partially impaired in Tlr2-deficient larvae at 6 and 12 hpi (Figures 3A and 3B). Moreover, Tlr2-deficient larvae also showed lower Nfkb activity than wild type larvae not only at the site of the injection but also in the head and the whole body at all analyzed timepoints (Figure 3C).

**Figure 3:**
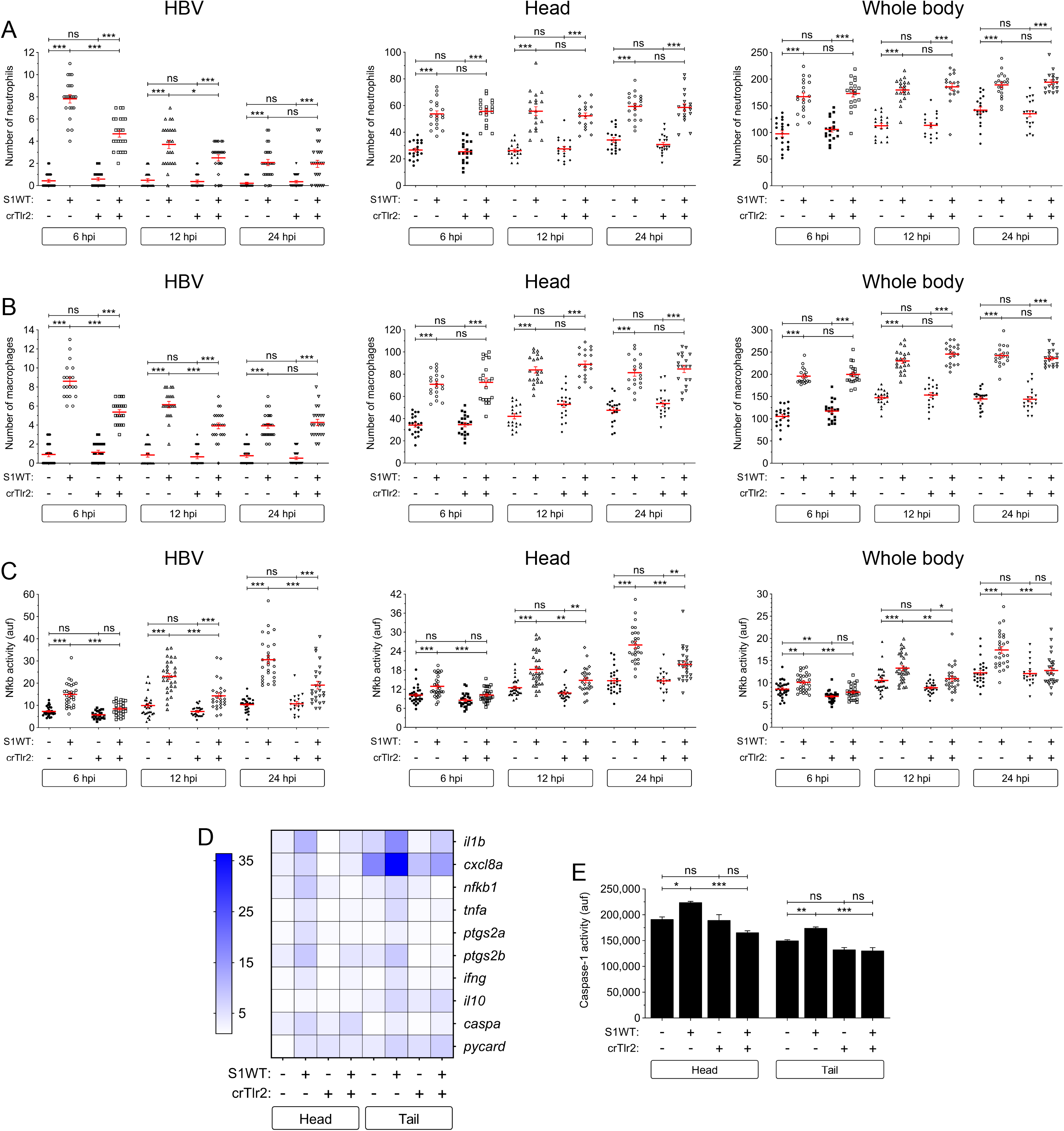
Tlr2 mediates the S1WT-induced hyperinflammation in zebrafish. One-cell stage zebrafish eggs of *Tg(lyz:dsRED2)* (A), *Tg(mfap4:mCherry)* (B), *Tg(NFkB-RE:eGFP)* (C) and wild type (D, E) were microinjected with control or *tlr2* crRNA/Cas9 complexes. At 2 dpf, recombinant S1WT (+) or vehicle (-) were injected in the hindbrain ventricle (HBV) of control and Tlr2-deficient larvae. Neutrophil (A) and macrophage (B) recruitment and number, and Nfkb activation (C) were analyzed at 6, 12 and 24 hpi by fluorescence microscopy, the transcript levels of the indicated genes were analyzed at 12 hpi by RT-qPCR (D), and caspase-1 activity was determined at 24 hpi using a fluorogenic substrate (E). Each dot represents one individual and the mean ± S.E.M. for each group is also shown. RT-qPCR data are depicted as a heat map in D with higher expression shown in darker color. P values were calculated using one-way ANOVA and Tukey multiple range test. ns, not significant, *≤p0.05, **p≤0.01, ***p≤0.001. auf, arbitrary units of fluorescence.

The above results were then confirmed by RT-qPCR. Thus, the transcript levels of *tlr2* significantly decreased in Tlr2-deficient animals, further confirming the high efficiency of the crRNA used (Figure S5A). Furthermore, the transcript levels of *il1b, cxcl8a, nfkb1, tnfa, ptgs2a, ptgs2b* and *ifng* were lower locally and systemically in the S1WT injected Tlr2-deficient larvae than in their wild type siblings (Figures 3D and S6A–S6G). However, no significant differences were observed in the mRNA levels of genes encoding anti-inflammatory I110 and the inflammasome components Caspa and Pycard (Figures 3H–3J). Surprisingly, the induction of caspase-1 by S1WT was also attenuated in Tlr2-deficient larvae locally and systemically (Figure 3E).

## DISCUSSION

Since the beginning of the COVID-19 pandemic, scientists are trying to find the molecular basis of SARS-CoV-2 virulence and host immune responses to find targeted treatments to moderate the severe symptoms of the disease and save people’s lives. Although both TLRs and the inflammasome pathways have been found to be involved in COVID-19, a mechanistic understanding of their involvement in COVID-19 progression is still unclear [25,26]. It has been suggested that the imbalance between the generation of excessive inflammation through TLR/MyD88 pathway and IFN-β/TRIF pathway plays a key role in COVID-19 severity [13]. Computer-based modelling has found that the S protein of SARS-CoV-2 is predicted to bind to TLR4 [15] and, more interestingly, SARS-CoV-2 S1 protein engaged TLR4 and strongly activated the inflammatory response leading to the production of pro-inflammatory mediators through nuclear factor κB (NF-κB) and stress-activated mitogen-activated protein kinase (MAPK) signaling pathways [14,27]. Moreover, it was suggested that SARS-CoV-2 S glycoprotein binds and activates TLR4, leading to increased cell surface expression of ACE2 which, in turn, would facilitate viral entry and cause the COVID-19-associated CSS [28]. Similarly, it has been proposed that the TLR2 signaling pathway is activated following SARS-CoV-2 infection, resulting in strong production of proinflammatory cytokines, suggesting that it may contribute to the severity of COVID-19 [29]. TLR2 is known to form heterodimers with TLR1 and TLR6, which increases its ligand diversity and allows detection of different kinds of pathogens, including viruses [11]. Recently, the envelope protein (E) of SARS-CoV-2 has been found to be a ligand of human and mouse TLR2 and, surprisingly, plays a critical role in COVID-19-associated CSS in the K18–hACE2 transgenic mice model [12]. However, this study has reported that S1+S2 of SARS-Cov-2 failed to activate mouse macrophages and human PBMCs [12]. In stark contrast, another study found that TLR2 recognizes the SARS-CoV-2 S protein and then dimerizes with TLR1 or TLR6 to activate the NF-κB pathway and promote CSS [30].

In the present study, we used a newly established zebrafish model of COVID-19 [24] based on the larval hindbrain injection of SARS-CoV-2 S1 protein to further understand the contribution of Tlr2 and Myd88 signaling pathway to the COVID-19-asscociated CSS. Genetic experiments demonstrated that Tlr2 sensed S1WT and induced hyperinflammation via Myd88 in zebrafish. Furthermore, this model has revealed: (i) S1WT-induced hyperinflammation is independent of the production of Il1b and (ii) S1WT-induced emergency myelopoiesis is Tlr2/Myd88-independent. On the one hand, the relevance of IL1B in the COVID-19-associated CSS is unclear, as a large controlled trial in hospitalized patients with COVID-19 found no therapeutic benefit of IL-1 blockade [31], whereas another trial of early treatment of COVID-19 patients with anakinra, to block IL-1, found decreased patient severity and improved survival [32]. On the other hand, although the relevance of emergency hematopoiesis in COVID-19 has been recognized, it has been less studied than the CSS. Thus, multi-omic single-cell immune profiling of COVID-19 patients has revealed that emergency myelopoiesis is a prominent feature of fatal COVID-19 [33]. In addition, COVID-19 patients in intensive care also show low levels of hemoglobin and circulating nucleated red cells, and erythroid progenitors can be infected by SARS-CoV-2 via ACE2 [34]. The zebrafish model is excellent for further understanding the role of altered hematopoiesis in patients with COVID-19. Thus, although our results point to the relevance of the inflammasome in S1-driven emergency myelopoiesis and dispensability of Tlr, the decreased caspase-1 activity levels in Tlr2- and Myd88-deficient larvae injected with S1 suggest a crosstalk between these 2 pivotal inflammatory pathways.

In summary, despite controversies, it appears that both TLR4 and TLR2 may contribute significantly to the pathogenesis of COVID-19 by promoting CSS. Therefore, TLR4 and TLR2 appear to be promising therapeutic targets in COVID-19 [35–37]. The zebrafish model of COVID-19 has confirmed the critical role played by TLR signaling pathway in COVID-19-associated CSS. This model, therefore, is an excellent platform for chemical screening of anti-inflammatory Tlr2 antagonist compounds to alleviate CSS and to identify therapeutic targets and novel drugs to treat COVID-19.

## MATERIALS AND METHODS

### Animals

Zebrafish (*Danio rerio* H.) were obtained from the Zebrafish International Resource Center and mated, staged, raised and processed as described [38]. The lines *Tg(mpx:eGFP)^i114^* [39], *Tg(lyz:DsRED2)^nz50^ [40], Tg(mfap4:mCherry-F)^ump6^* referred to as *Tg(mfap4:mCherry)* [41], *Tg(NFkB-RE:eGFP)^sh235^* referred to as *nfkb:eGFP* [42], *myd88^hu3668/hu3568^* mutant [43], and casper (*mitfa^w2/w2^; mpv17^a9/a9^)* [44] were previously described. The experiments performed comply with the Guidelines of the European Union Council (Directive 2010/63/EU) and the Spanish RD 53/2013. The experiments and procedures were performed approved by the Bioethical Committees of the University of Murcia (approval number #669/2020).

### Analysis of gene expression

Total RNA was extracted from whole head/tail part of the zebrafish body with TRIzol reagent (Invitrogen) following the manufacturer’s instructions and treated with DNase I, amplification grade (1 U/mg RNA: Invitrogen). SuperScript IV RNase H Reverse Transcriptase (Invitrogen) was used to synthesize first-strand cDNA with random primer from 1mg of total RNA at 50 °C for 50 min. Real-time PCR was performed with an ABIPRISM 7500 instrument (Applied Biosystems) using SYBR Green PCR Core Reagents (Applied Biosystems). The reaction mixtures were incubated for 10 min at 95 °C, followed by 40 cycles of 15 s at 95 °C, 1 min at 60 °C, and finally 15 s at 95 °C, 1 min 60 °C, and 15 s at 95 °C. For each mRNA, gene expression was normalized to the ribosomal protein S11 (rps11) content in each sample using the Pfaffl method [45]. The primers used are shown in Table S1. In all cases, each PCR was performed with samples in triplicate and repeated with at least two independent samples.

### CRISPR and recombinant protein injections, and chemical treatments in zebrafish

crRNA for zebrafish *il1b, tlr2* (Table S2) or negative control (Catalog #1072544), and tracrRNA were resuspended in Nuclease-Free Duplex Buffer to 100 μM. One μl of each was mixed and incubated for 5 min at 95 °C for duplexing. After removing from the heat and cooling to room temperature, 1.43 μl of Nuclease-Free Duplex Buffer was added to the duplex, giving a final concentration of 1000 ng/μl. Finally, the injection mixture was prepared by mixing 1 μl of duplex, 2.55 μl of Nuclease-Free Duplex Buffer, 0.25 μl Cas9 Nuclease V3 (IDT, 1081058) and 0.25 μl of phenol red, resulting in final concentrations of 250 ng/μl of gRNA duplex and 500 ng/μl of Cas9. The prepared mix was microinjected into the yolk sac of one- to eight-cell-stage embryos using a microinjector (Narishige) (0.5-1 nl per embryo). The same amounts of gRNA were used in all experimental groups. The efficiency of gRNA was checked by amplifying the target sequence with a specific pair of primers (Table S1) and the TIDE webtool (https://tide.nki.nl/) and/or SYNTHEGO Crisper Performance Analysis webtool (https://ice.synthego.com). Embryos injected with crIl1b or crTlr2 were sorted at 2 hpf to choose the ones in the same developmental stage and raised at similar densities. At 24 hpf, the number of dead/alive embryos was determined and within the surviving group the number of embryos with any malformation was scored. At 26 hpf, the number of otic vesicle structures that could fit between the eye and otic vesicle in each larva were estimated. The higher the number of otic vesicles fitted, the lower the level of the larval development [46].

Recombinant His-tagged Spike S1 wild-type produced in baculovirus-insect cells and with < 1.0 EU per μg protein as determined by the LAL method (#40591-V08B1, Sino Biological) or flagellin (Invivogen) at a concentration of 0.25 mg/ml supplemented with phenol red were injected into the hindbrain ventricle (1 nl) of 48 hpf zebrafish larvae.

In some experiments, 24hpf embryos were treated with 0.3% N-Phenylthiourea (PTU) to inhibit melanogenesis.

### Caspase-1 activity assays

Caspase-1 activity was determined with the fluorometric substrate Z-YVAD 7-Amido-4-trifluoromethylcoumarin (Z-YVAD-AFC, caspase-1 substrate VI, Calbiochem) as described previously [47,48]. A representative graph of caspase-1 activity of three repeats is shown in the figures.

### In vivo imaging

To study immune cell recruitment to the injection site and Nfkb activation, 2 dpf *mpx:eGFP, mfap4:mcherry* or *nfkb:egfp* larvae were anaesthetized in embryonic medium with 0.16 mg/ml buffered tricaine. Images of the hindbrain, head or whole-body area were taken 3, 6, 12 and 24 h post-injection (hpi) using a Leica MZ16F fluorescence stereomicroscope. The number of neutrophils or macrophages was determined by visual counting and the fluorescence intensity was obtained and analyzed with ImageJ (FIJI) software [49].

Neutral red stains zebrafish macrophage granules and the procedure was performed as originally reported [50]. Briefly, macrophage staining was performed on live 3 dpf larvae and was obtained by incubating the embryos in 2.5 g/ml of neutral red (in embryonic medium) at 25-30°C in the dark for 5-8 h. The larvae were anesthetized in 0.16 mg/ml buffered tricaine and imaged using a Leica MZ16F fluorescence stereo microscope.

In all experiments, images were pooled from at least 3 independent experiments performed by two people and using blinded samples.

### Statistical analysis

Data are shown as mean ± s.e.m. and were analyzed by analysis of variance and a Tukey multiple range test to determine differences between groups. The differences between two samples were analyzed by Student’s t-test.

## ACKNOWLEDGMENTS

We thank I. Fuentes and P. Martinez for their excellent technical assistance, and Profs. Tobin, Crosier, Renshaw, Zon, Meijer and Lutfalla for the zebrafish lines.

## AUTHOR CONTRIBUTIONS

SDT, VM and MLC conceived the study; SDT, AM-L, AP and SC performed the research; SDT, AM-L, AP, SC, MLC and VM analyzed the data; and SDT and VM wrote the manuscript with minor contributions from other authors.

## DATA AVAILABILITY STATEMENT

All data needed to evaluate the conclusions in the paper are present in the paper and/or the Supplementary Materials.

---

**Figure S1.**
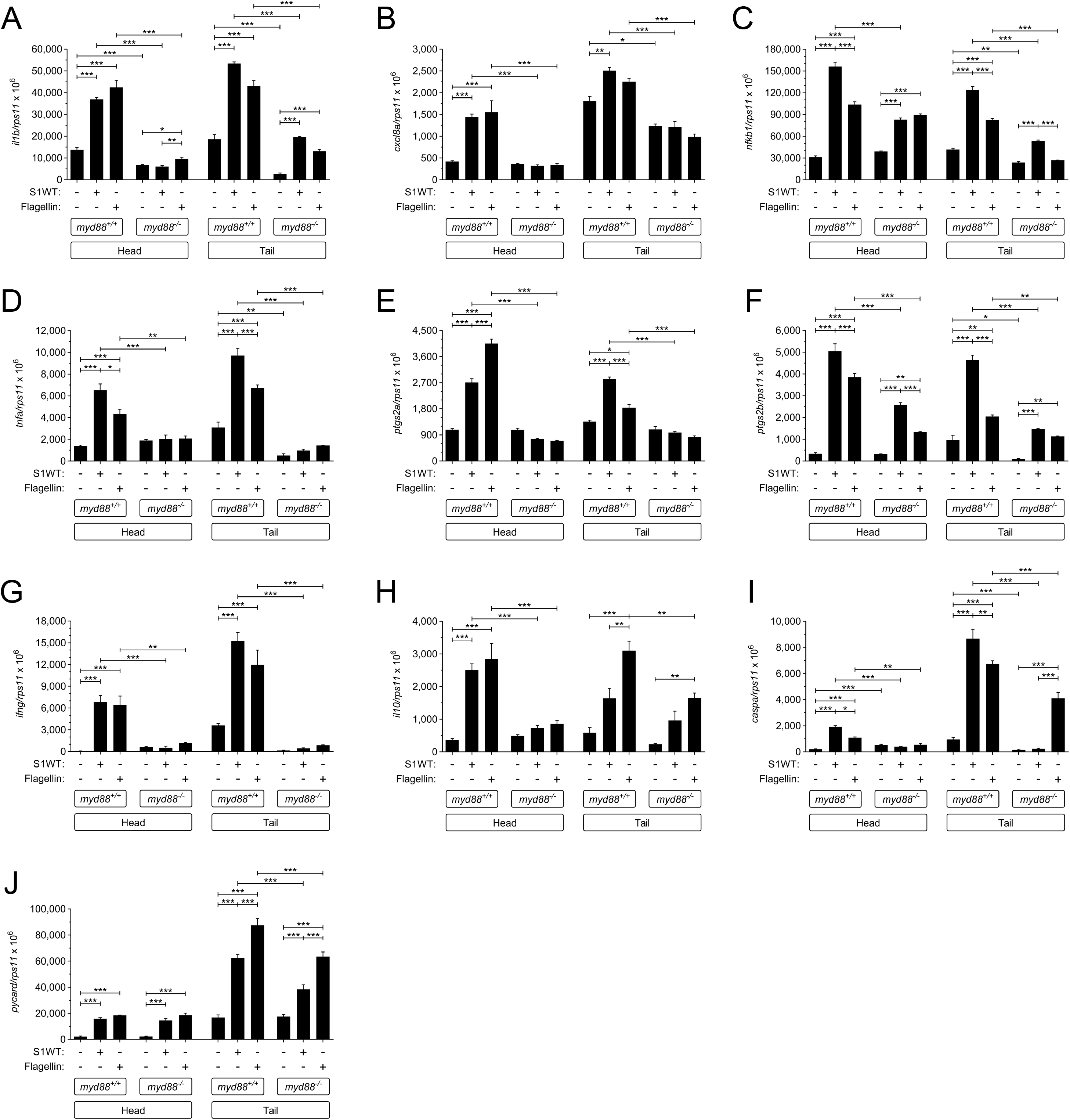
(related to Figure 1). Gene expression analysis of wild type and Myd88-deficient larvae injected with wild type S1. Recombinant S1WT (+) or vehicle (-) were injected in the hindbrain ventricle of 2 dpf wild type and Myd88-deficient larvae, and the transcript levels of the indicated genes were analyzed at 12 hpi by RT-qPCR in larval head and tail. Data are shown as mean ± S.E.M. P values were calculated using one way ANOVA and Tukey multiple range test. ns, not significant, *≤p0.05, **p≤0.01, ***p≤0.001.

**Figure S2.**
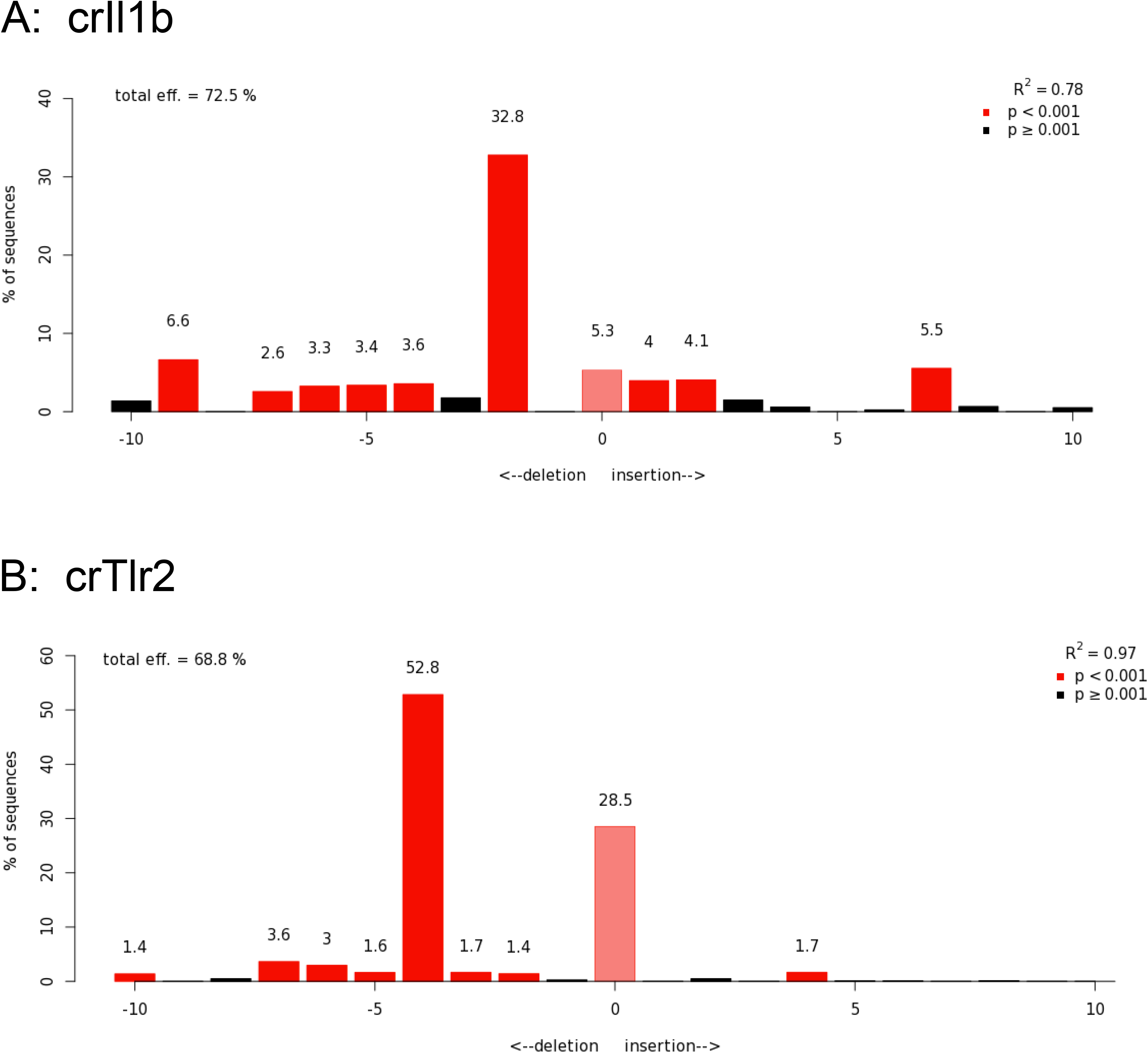
(related to Figures 2 and 3). Analysis of genome editing efficiency in larvae injected with *il1b* (A) and *tlr2* (B) crRNA/Cas 9 complexes and quantification rate of nonhomologous end joining mediated repair showing all insertions and deletions (INDELS) at the target site using TIDE (https://tide.nki.nl).

**Figure S3.**
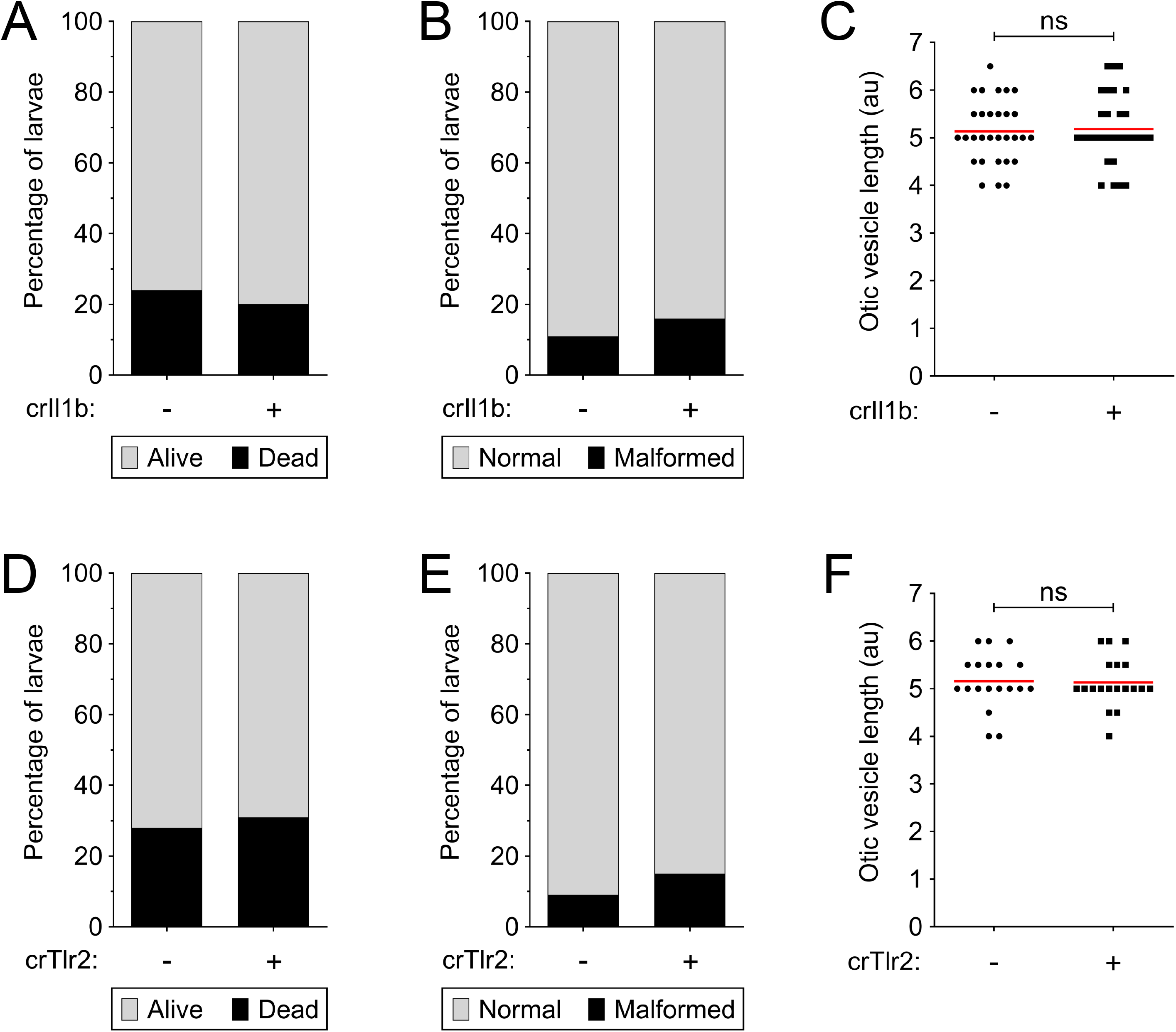
(related to Figures 2 and 3). Il1b and Tlr2 deficiencies do not affect larval development. The precentage of dead/alive (A, D) and malformed embryos (B, E), and the developmental stage (C, F) of Il1b- (A-C) and Tlr2-deficient (D-F) embryos was determined at 24 hpf. au, arbitrary units.

**Figure S4.**
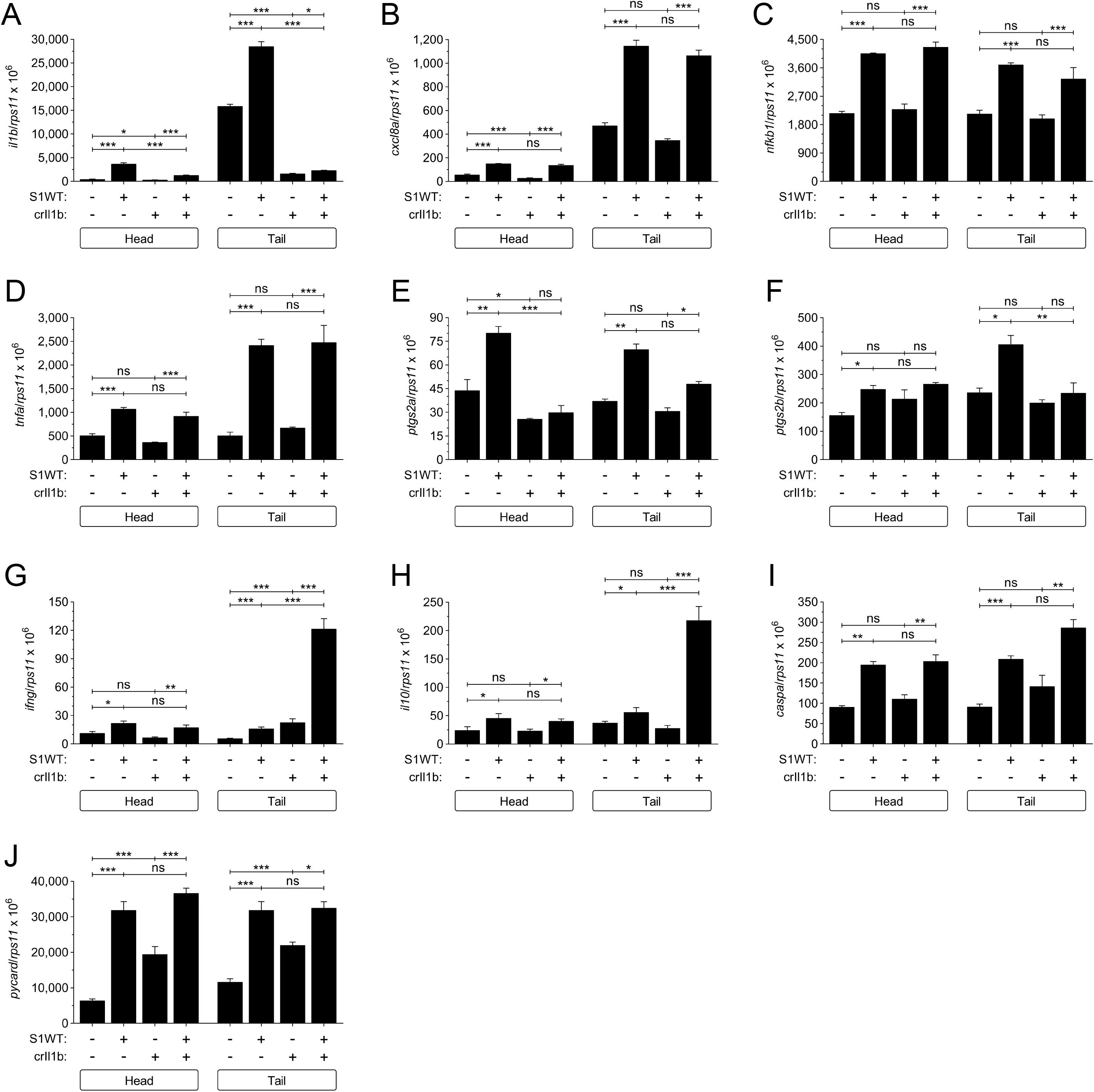
(related to Figure 2). Gene expression analysis of wild type and Il1b-deficient larvae injected with wild type S1. Recombinant S1WT (+) or vehicle (-) were injected in the hindbrain ventricle of 2 dpf wild type and Il1b-deficient larvae, and the transcript levels of the indicated genes were analyzed at 12 hpi by RT-qPCR in larval head and tail. Data are shown as mean ± S.E.M. P values were calculated using one way ANOVA and Tukey multiple range test. ns, not significant, *≤p0.05, **p≤0.01, ***p≤0.001.

**Figure S5.**
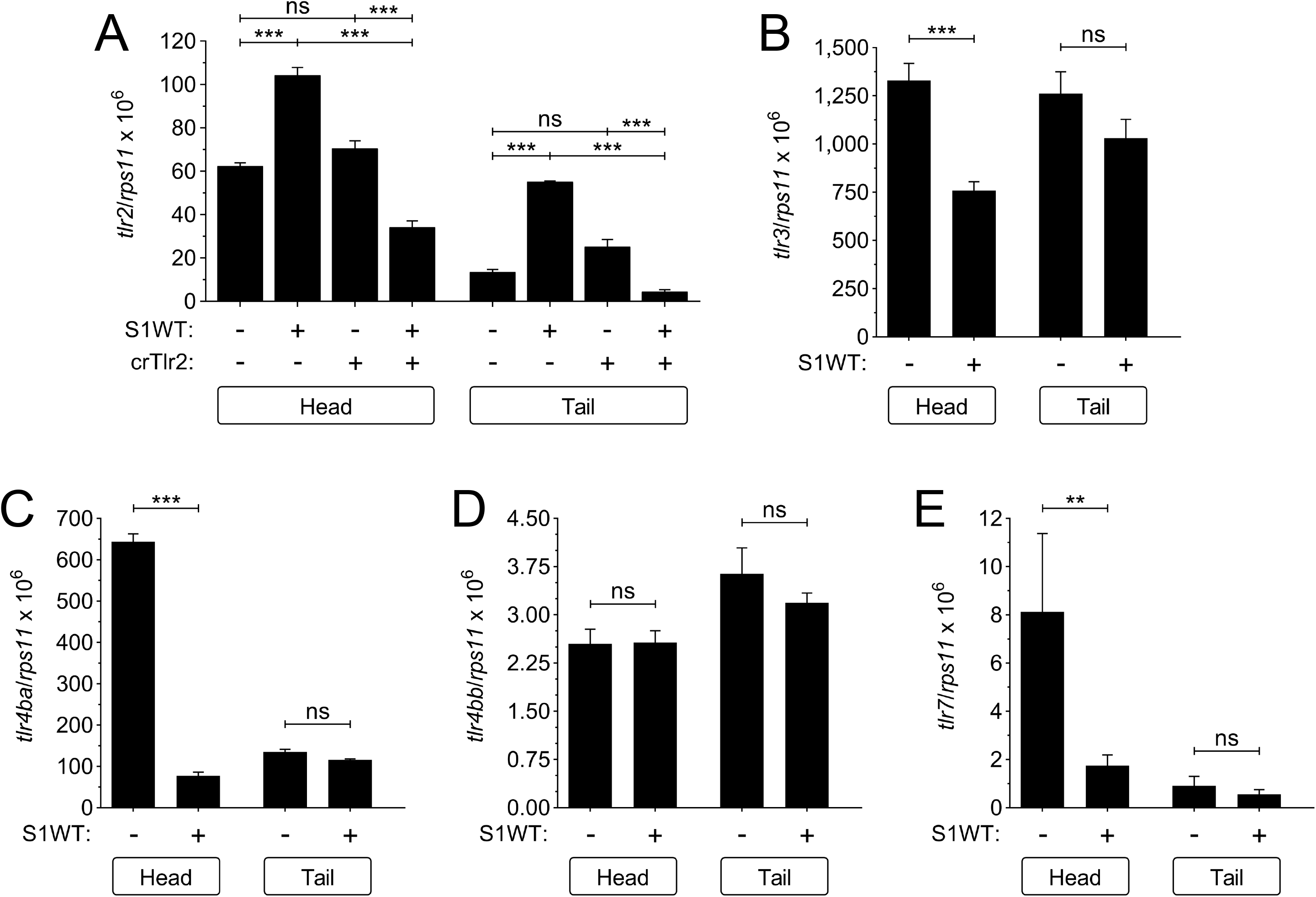
(related to Figure 3). Tlr expression analysis of wild type and Tlr2-deficient larvae injected with wild type S1. Recombinant S1WT (+) or vehicle (-) were injected in the hindbrain ventricle of 2 dpf wild type and Tlr2-deficient larvae, and the transcript levels of the indicated *tlr* genes were analyzed at 12 hpi by RT-qPCR in larval head and tail. Data are shown as mean ± S.E.M. P values were calculated using one way ANOVA and Tukey multiple range test. ns, not significant, **p≤0.01, ***p≤0.001.

**Figure S6.**
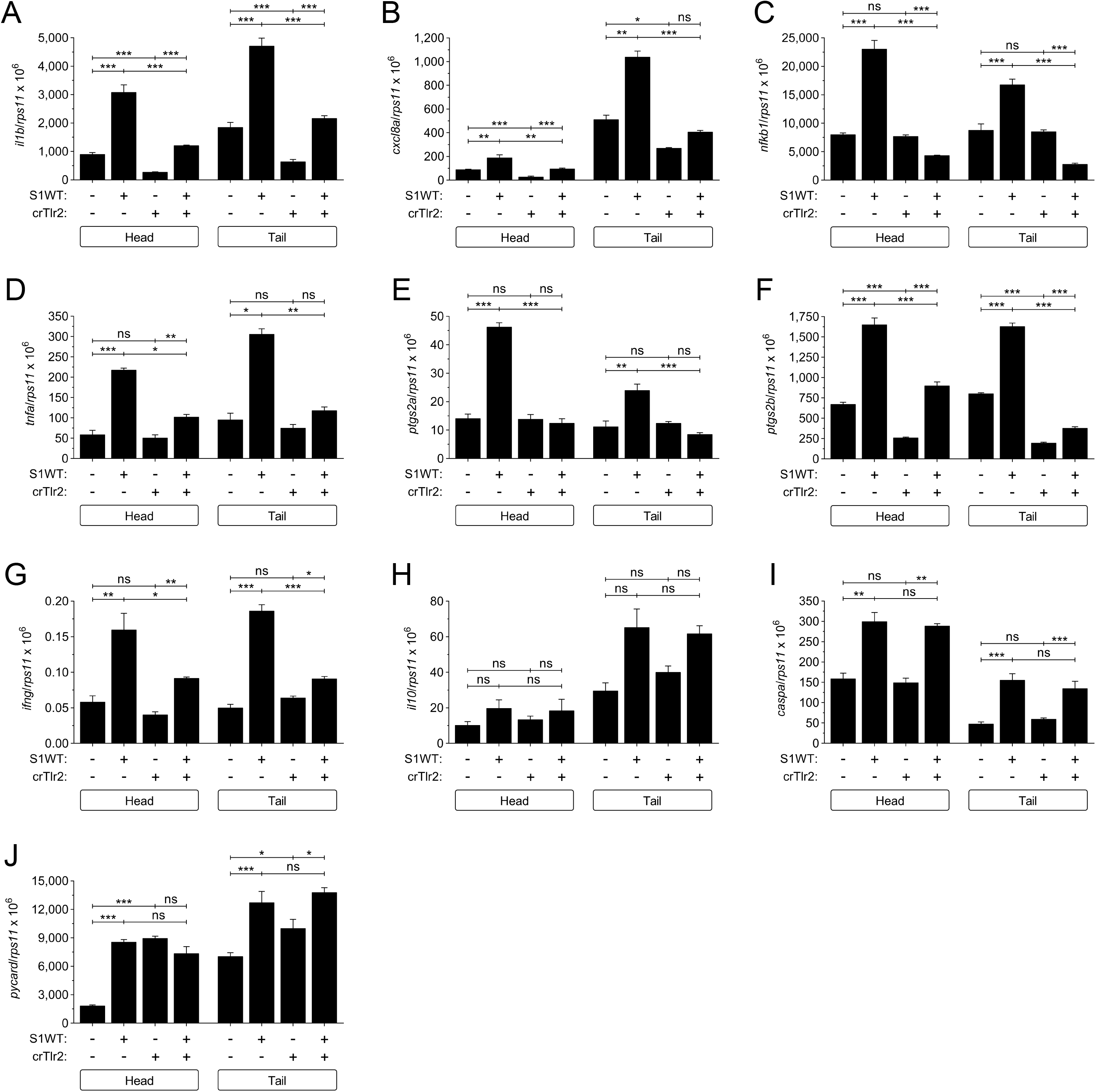
(related to Figure 3). Gene expression analysis of wild type and Tlr2-deficient larvae injected with wild type S1. Recombinant S1WT (+) or vehicle (-) were injected in the hindbrain ventricle of 2 dpf wild type and Tlr2-deficient larvae, and the transcript levels of the indicated genes were analyzed at 12 hpi by RT-qPCR in larval head and tail. Data are shown as mean ± S.E.M. P values were calculated using one way ANOVA and Tukey multiple range test. ns, not significant, *≤p0.05, **p≤0.01, ***p≤0.001.

**Table S1.**
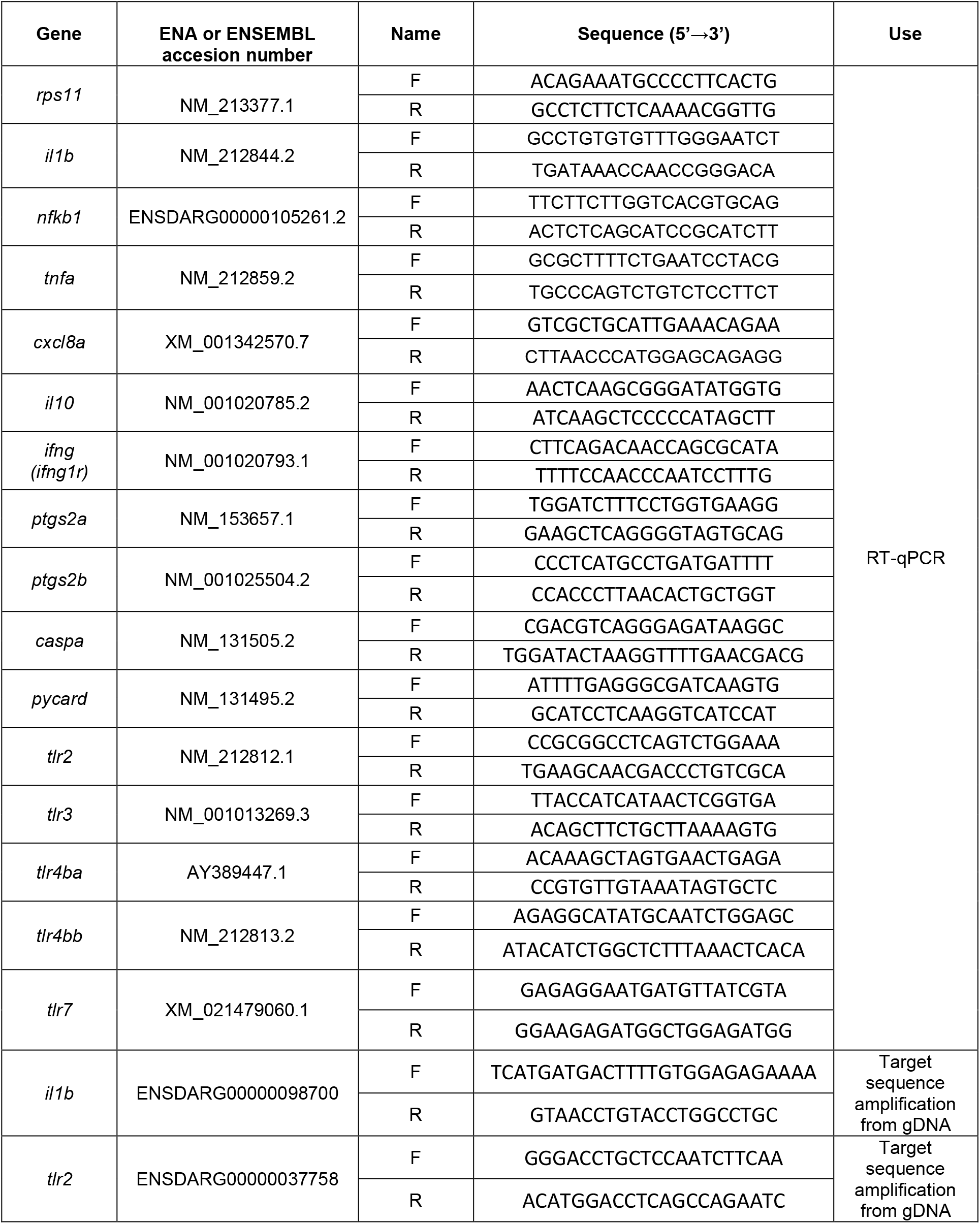
Primers used in this study. The gene symbols followed the Zebrafish Nomenclature Guidelines (http://zfin.org/zf_info/nomen.html). Ena, European Nucleotide Archive.

**Table S2.**
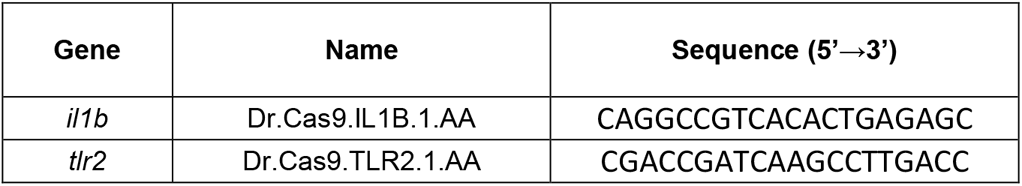
crRNA used in this study. The gene symbols followed the Zebrafish Nomenclature Guidelines (http://zfin.org/zf_info/nomen.html).

## References

1. Lester SN, Li K. Toll-like receptors in antiviral innate immunity. J Mol Biol. 2014 Mar 20;426(6):1246–64.

2. Kawai T, Akira S. Toll-like receptors and their crosstalk with other innate receptors in infection and immunity. Immunity. 2011 May 27;34(5):637–50.

3. Akira S, Takeda K. Toll-like receptor signalling. Nat Rev Immunol. 2004 Jul;4(7):499–511.

4. Akira S. TLR signaling. Curr Top Microbiol Immunol. 2006;311:1–16.

5. Kawai T, Akira S. The role of pattern-recognition receptors in innate immunity: update on Toll-like receptors. Nat Immunol. 2010 May;11(5):373–84.

6. Yamamoto M, Sato S, Mori K, et al. Cutting edge: a novel Toll/IL-1 receptor domain-containing adapter that preferentially activates the IFN-beta promoter in the Toll-like receptor signaling. J Immunol. 2002 Dec 15;169(12):6668–72.

7. Takeda K, Kaisho T, Akira S. Toll-like receptors. Annu Rev Immunol. 2003;21:335–76.

8. Bieback K, Lien E, Klagge IM, et al. Hemagglutinin protein of wild-type measles virus activates toll-like receptor 2 signaling. J Virol. 2002 Sep;76(17):8729–36.

9. Kurt-Jones EA, Popova L, Kwinn L, et al. Pattern recognition receptors TLR4 and CD14 mediate response to respiratory syncytial virus. Nat Immunol. 2000 Nov;1(5):398–401.

10. Shuang Chen, Wong MH, Schulte DJ, et al. Differential expression of Toll-like receptor 2 (TLR2) and responses to TLR2 ligands between human and murine vascular endothelial cells. J Endotoxin Res. 2007;13(5):281–96.

11. Oliveira-Nascimento L, Massari P, Wetzler LM. The Role of TLR2 in Infection and Immunity. Front Immunol. 2012;3:79.

12. Zheng M, Karki R, Williams EP, et al. TLR2 senses the SARS-CoV-2 envelope protein to produce inflammatory cytokines. Nat Immunol. 2021 Jul;22(7):829–838.

13. Mabrey FL, Morrell ED, Wurfel MM. TLRs in COVID-19: How they drive immunopathology and the rationale for modulation. Innate Immun. 2021 10;27(7-8):503–513.

14. Zhao Y, Kuang M, Li J, et al. SARS-CoV-2 spike protein interacts with and activates TLR41. Cell Res. 2021 Jul;31(7):818–820.

15. Choudhury A, Mukherjee S. In silico studies on the comparative characterization of the interactions of SARS-CoV-2 spike glycoprotein with ACE-2 receptor homologs and human TLRs. J Med Virol. 2020 10;92(10):2105–2113.

16. Zhang J, Kong X, Zhou C, et al. Toll-like receptor recognition of bacteria in fish: ligand specificity and signal pathways. Fish Shellfish Immunol. 2014 Dec;41(2):380–8.

17. Jault C, Pichon L, Chluba J. Toll-like receptor gene family and TIR-domain adapters in Danio rerio. Mol Immunol. 2004 Jan;40(11):759–71.

18. Meijer AH, Gabby Krens SF, Medina Rodriguez IA, et al. Expression analysis of the Toll-like receptor and TIR domain adaptor families of zebrafish. Mol Immunol. 2004 Jan;40(11):773–83.

19. Matsuo A, Oshiumi H, Tsujita T, et al. Teleost TLR22 recognizes RNA duplex to induce IFN and protect cells from birnaviruses. J Immunol. 2008 Sep 1;181(5):3474–85.

20. Yeh DW, Liu YL, Lo YC, et al. Toll-like receptor 9 and 21 have different ligand recognition profiles and cooperatively mediate activity of CpG-oligodeoxynucleotides in zebrafish. Proc Natl Acad Sci U S A. 2013 Dec 17;110(51):20711–6.

21. Li Y, Cao X, Jin X, et al. Pattern recognition receptors in zebrafish provide functional and evolutionary insight into innate immune signaling pathways. Cell Mol Immunol. 2017 01;14(1):80–89.

22. Yang S, Marin-Juez R, Meijer AH, et al. Common and specific downstream signaling targets controlled by Tlr2 and Tlr5 innate immune signaling in zebrafish. BMC Genomics. 2015 Jul 25;16:547.

23. Stein C, Caccamo M, Laird G, et al. Conservation and divergence of gene families encoding components of innate immune response systems in zebrafish. Genome Biol. 2007;8(11):R251.

24. Tyrkalska SD, Martínez-López A, Arroyo AB, et al. A zebrafish model of COVID-19-associated cytokine storm syndrome reveals differential proinflammatory activities of Spike proteins of SARS-CoV-2 variants of concern. bioRxiv. 2021:2021.12.05.471277.

25. Vora SM, Lieberman J, Wu H. Inflammasome activation at the crux of severe COVID-19. Nat Rev Immunol. 2021 Aug 09.

26. Khanmohammadi S, Rezaei N. Role of Toll-like receptors in the pathogenesis of COVID-19. J Med Virol. 2021 May;93(5):2735–2739.

27. Shirato K, Kizaki T. SARS-CoV-2 spike protein S1 subunit induces pro-inflammatory responses via toll-like receptor 4 signaling in murine and human macrophages. Heliyon. 2021 Feb;7(2):e06187.

28. Aboudounya MM, Heads RJ. COVID-19 and Toll-Like Receptor 4 (TLR4): SARS-CoV-2 May Bind and Activate TLR4 to Increase ACE2 Expression, Facilitating Entry and Causing Hyperinflammation. Mediators Inflamm. 2021;2021:8874339.

29. Sariol A, Perlman S. SARS-CoV-2 takes its Toll. Nat Immunol. 2021 07;22(7):801–802.

30. Khan S, Shafiei MS, Longoria C, et al. SARS-CoV-2 spike protein induces inflammation via TLR2-dependent activation of the NF-κB pathway. Elife. 2021 12 06;10.

31. Declercq J, Van Damme KFA, De Leeuw E, et al. Effect of anti-interleukin drugs in patients with COVID-19 and signs of cytokine release syndrome (COV-AID): a factorial, randomised, controlled trial. Lancet Respir Med. 2021 Dec;9(12):1427–1438.

32. Kyriazopoulou E, Poulakou G, Milionis H, et al. Early treatment of COVID-19 with anakinra guided by soluble urokinase plasminogen receptor plasma levels: a double-blind, randomized controlled phase 3 trial. Nat Med. 2021 Oct;27(10):1752–1760.

33. Wilk AJ, Lee MJ, Wei B, et al. Multi-omic profiling reveals widespread dysregulation of innate immunity and hematopoiesis in COVID-19. J Exp Med. 2021 Aug 2;218(8).

34. Huerga Encabo H, Grey W, Garcia-Albornoz M, et al. Human Erythroid Progenitors Are Directly Infected by SARS-CoV-2: Implications for Emerging Erythropoiesis in Severe COVID-19 Patients. Stem Cell Reports. 2021 Mar 9;16(3):428–436.

35. Voelker DR, Numata M. Phospholipid regulation of innate immunity and respiratory viral infection. J Biol Chem. 2019 03 22;294(12):4282–4289.

36. Durai P, Shin HJ, Achek A, et al. Toll-like receptor 2 antagonists identified through virtual screening and experimental validation. FEBS J. 2017 07;284(14):2264–2283.

37. Proud PC, Tsitoura D, Watson RJ, et al. Prophylactic intranasal administration of a TLR2/6 agonist reduces upper respiratory tract viral shedding in a SARS-CoV-2 challenge ferret model. EBioMedicine. 2021 Jan;63:103153.

38. Westerfield M. The Zebrafish Book. A Guide for the Laboratory Use of Zebrafish Danio* (Brachydanio) rerio.. Eugene, OR.: University of Oregon Press.; 2000.

39. Renshaw SA, Loynes CA, Trushell DM, et al. A transgenic zebrafish model of neutrophilic inflammation. Blood. 2006 Dec 15;108(13):3976–8.

40. Hall C, Flores MV, Storm T, et al. The zebrafish lysozyme C promoter drives myeloid-specific expression in transgenic fish. BMC Dev Biol. 2007;7:42.

41. Phan QT, Sipka T, Gonzalez C, et al. Neutrophils use superoxide to control bacterial infection at a distance. PLoS Pathog. 2018 Jul;14(7):e1007157.

42. Kanther M, Sun X, Mühlbauer M, et al. Microbial colonization induces dynamic temporal and spatial patterns of NF-κB activation in the zebrafish digestive tract. Gastroenterology. 2011 Jul;141(1):197–207.

43. van der Vaart M, Spaink HP, Meijer AH. Pathogen recognition and activation of the innate immune response in zebrafish. Adv Hematol. 2012;2012:159807.

44. White RM, Sessa A, Burke C, et al. Transparent adult zebrafish as a tool for in vivo transplantation analysis. Cell Stem Cell. 2008 Feb 07;2(2): 183–9.

45. Pfaffl MW. A new mathematical model for relative quantification in real-time RT-PCR. Nucleic Acids Res. 2001 May 1;29(9):e45.

46. Kimmel CB, Ballard WW, Kimmel SR, et al. Stages of embryonic development of the zebrafish. Dev Dyn. 1995 Jul;203(3):253–310.

47. Lopez-Castejon G, Sepulcre MP, Mulero I, et al. Molecular and functional characterization of gilthead seabream Sparus aurata caspase-1: the first identification of an inflammatory caspase in fish. Mol Immunol. 2008 Jan;45(1):49–57.

48. Angosto D, López-Castejón G, López-Muñoz A, et al. Evolution of inflammasome functions in vertebrates: Inflammasome and caspase-1 trigger fish macrophage cell death but are dispensable for the processing of IL-1β. Innate Immun. 2012 Dec;18(6):815–24.

49. Schindelin J, Arganda-Carreras I, Frise E, et al. Fiji: an open-source platform for biological-image analysis. Nat Methods. 2012 Jun 28;9(7):676–82.

50. Herbomel P, Thisse B, Thisse C. Zebrafish early macrophages colonize cephalic mesenchyme and developing brain, retina, and epidermis through a M-CSF receptor-dependent invasive process. Dev Biol. 2001 Oct 15;238(2):274–88.

